# FOXM1 Expression in Invasive Ductal Breast Carcinoma of No Special Type: Insights from RNA-seq and Immunohistochemical Analysis

**DOI:** 10.1101/2025.09.07.674683

**Authors:** Masahiro Ohara, Emi Mikami, Wakako Inohana, Taeko Kurosawa, Ayako Nakame, Ayaka Sakakibara, Yuki Ichinose, Akihiro Fujimoto, Asami Nukui, Aya Asano, Hiroko Shimada, Kyoko Asai, Masataka Hirasaki, Hideki Yokogawa, Kazuo Matsuura, Hiroshi Ishiguro, Takahiro Hasebe, Nobuko Fujiuchi, Akihiko Osaki, Toshiaki Saeki

## Abstract

**Background:** Invasive ductal carcinoma of no special type (IDC-NST) is the most common subtype of breast cancer, characterized by significant clinical heterogeneity. Forkhead box M1 (FOXM1) is a key transcription factor involved in cell cycle regulation and tumor progression, but its expression profile and clinical significance in IDC-NST remain incompletely understood.

**Methods:** We analyzed FOXM1 expression in a cohort of 100 IDC-NST patients using RNA sequencing and immunohistochemistry (IHC). FOXM1 mRNA levels were quantified, and protein expression was scored based on the percentage of positive tumor cells. Differentially expressed genes (DEGs) between high and low FOXM1 expression groups were identified, followed by pathway enrichment and protein–protein interaction (PPI) network analyses. The prognostic value of FOXM1 was evaluated by recurrence-free survival (RFS) analysis.

**Results:** FOXM1 protein expression correlated significantly with mRNA abundance in 22 representative cases (p < 0.05). Receiver operating characteristic (ROC) curve analysis identified a cut-off value of 4.954 CPM for FOXM1 mRNA to predict recurrence (AUC = 0.642). High FOXM1 expression was associated with larger tumor size, higher histological grade, and negative hormone receptor status. Patients with high FOXM1 expression exhibited significantly poorer 5-year RFS (73.6% vs. 92.3%, p < 0.001). Multivariate analysis confirmed FOXM1 as an independent prognostic factor (HR 15.26; p = 0.026). Transcriptomic profiling revealed 190 upregulated and 197 downregulated genes in the FOXM1-high group, enriched in cell cycle and mitotic pathways. PPI network analysis positioned FOXM1 as a central hub coordinating genes involved in chromosomal stability and mitosis.

**Conclusions:** FOXM1 overexpression is a strong independent predictor of poor prognosis in IDC-NST and plays a critical role in tumor proliferation and genome integrity. These findings support FOXM1 as a potential prognostic biomarker and therapeutic target, warranting further investigation into FOXM1-targeted therapies for breast cancer management.

## Introduction

Breast cancer remains the most commonly diagnosed malignancy and a leading cause of cancer-related death among women worldwide (1). Among its histological subtypes, invasive ductal carcinoma of no special type (IDC-NST) represents the most prevalent form, accounting for approximately 70–80% of all breast cancer cases (2,3). Despite advances in early detection and molecular subtyping, IDC-NST continues to exhibit variable clinical outcomes due to its heterogeneous nature (4). Understanding the molecular drivers that underpin this heterogeneity is critical for identifying novel prognostic biomarkers and therapeutic targets (5,6).

Forkhead box M1 (FOXM1), a transcription factor belonging to the FOX family, plays a central role in cell cycle regulation, DNA damage repair, and tumor progression (7,8). Aberrant expression of FOXM1 has been reported in a wide range of human malignancies, including breast cancer, where it is associated with increased tumor proliferation, genomic instability, and resistance to therapy (9–11). However, the precise expression profile of FOXM1 in IDC-NST and its potential clinical relevance remain insufficiently characterized, particularly when integrating transcriptomic and protein-level data.

In this study, we aimed to evaluate the expression of FOXM1 in IDC-NST by combining RNA sequencing (RNA-seq) and immunohistochemical (IHC) analyses. Importantly, all data were derived from our original patient cohort, without the use of any publicly available databases. By correlating transcriptomic data with tissue-based protein expression, we sought to provide comprehensive insights into the role of FOXM1 in this breast cancer subtype. This integrative approach may reveal novel associations between FOXM1 expression patterns and clinicopathological features, offering potential implications for prognostication and personalized therapeutic strategies.

## Materials and methods

### Patient and histological examinations

The study cohort consisted of 100 patients with invasive breast carcinoma of no special type (IBC-NST) who underwent surgical resection without prior neoadjuvant therapy at the Saitama Medical University International Medical Center between April 2007 and December 2015. During this period, a total of 855 patients underwent surgery for primary breast cancer at our institution. Of these, 100 cases with sufficient tumor tissue and RNA quality were selected for RNA sequencing analysis. This cohort included 26 cases previously reported in Ichinose et al. (2023) (12), and an additional 74 cases selected from earlier cases in the same period.

All the tumors were classified according to the pathological Union for International Cancer Control (UICC) TNM (pTNM) classification(13). The protocol for this study was approved by the institutional review board of the Saitama Medical University International Medical Center (IRB protocol number: # 2023-105). For histopathological examination of the tumors, the surgically resected specimens were fixed in 10% formalin. We defined the estrogen receptor status and progesterone receptor status of the tumor cells according to the American Society of Oncology/College of American Pathologists (ASCO/CAP) guideline (14). Human epidermal growth factor 2 (HER2) expression in the tumor cells was also categorized according to the ASCO/CAP guideline (15–17).

### RNA extraction

Total RNA isolation and library preparation were performed as previously reported (Ichinose et al., 2023)(1). Briefly, total RNA was extracted from FFPE IBC-NST specimens using the Maxwell® RSC RNA FFPE Kit (Promega). RNA quality was assessed by DV200 score with the Agilent 2200 TapeStation. Atier ribosomal RNA depletion with the Ribo-Zero Gold Kit (Illumina), libraries were prepared using the NEBNext Ultra RNA Library Prep Kit for Illumina (NEB) and sequenced on the Illumina HiSeqX platiorm (2 × 150-bp paired-end reads).

### Immunohistochemistry

Atier sectioning, paraffin-embedded tissue sections were deparaffinized and antigen retrieval was accomplished using high pH conditional buffer. Sections were incubated with anti-FOXM1 antibody overnight at 4℃. On the following day, sections were incubated for 30 min in Histofine Simple Stain MAX PO(MULTI) (Nichirei Bioscience,Tokyo,Japan), followed by incubating for 5 min in 3,3’-Diaminobenzidine (DAB) (Nichirei Bioscience). Sections were counterstained with hematoxylin. The percentage of positive cells was determined by counting at least 200 cells at hot spot areas for each specimen. Four staining categories were established—Tumor cells were scored as follows: 0 (no detectable staining), 1+ (positive staining in ≤10% of tumor cells), 2+ (positive staining in (>10% and ≤50%) of tumor cells, and 3+ (positive staining in >50% of tumor cells).

### Analysis of differentially expressed genes (DEGs)

Sequencing reads were first mapped to the rRNA reference (Human_rRNA_Reference_U13369) using Bowtie to remove rRNA reads. Unpaired reads were then filtered out using *fastq pair*. The remaining paired-end reads were aligned to the GRCh38 reference genome with STAR v2.7.8a. Transcript quantification was performed using RSEM to obtain expected counts and TPM (transcripts per kilobase million) values. TPM values were log₂-transformed atier adding +1 for downstream visualization and clustering analyses.

For differential expression analysis, the expected counts estimated by RSEM were analyzed using the *edgeR* package in R. Counts were normalized to CPM (counts per million) to account for library size differences. Statistical significance, including p-values and false discovery rates (FDR), was determined based on the normalized count data. DEGs were defined as those showing at least a twofold change in expression between the FOXM1-high and FOXM1-low groups, with p < 0.05.

The cutoff value for FOXM1 expression (4.954 CPM, based on normalized counts in edgeR) was determined from recurrence-free survival analysis. The cohort was then divided into two groups: FOXM1 > 4.954 CPM (n = 41) and FOXM1 < 4.954 CPM (n = 59). For enrichment analysis, DAVID (18) was used to perform Kyoto Encyclopedia of Genes and Genomes (KEGG) (19) pathway and Gene Ontology (GO) (20) analyses.

### STRING protein network analysis

To construct a protein–protein interaction (PPI) network, the STRING database (https://string-db.org/) was used with the cut-off criterion of a combined score > 0.7.

## Results

### Patient and tumor characteristics

Of 855 primary invasive ductal carcinoma cases treated with upfront surgery between April 2007 and December 2015, RNA was extracted from 100 cases. RNA-seq was performed on 74 newly selected cases, starting with the oldest specimens, in addition to the 26 cases previously reported (Ichinose et al., 2023). Table 1 shows the clinicopathological characteristics of the 100 patients. All the patients were Japanese women, ranging in age from 27 to 86 years (median, 59 years). The majority of the patients had pathological T1-stage tumors, 26 (26.0%) had lymph node involvement, 39 (39.0%) of the tumors were positive for lymphovascular invasion. A total of 69 (69.0%) tumors were ER-positive and 49 (49.0%) were PgR-positive according to IHC, and HER2-positive status was defined according to ASCO/CAP guidelines as either an IHC score of 3+ or an IHC score of 2+ with confirmation of gene amplification by FISH (HER2/CEP17 ratio ≥2.0); 20 cases (20.0%) were classified as HER2-positive by these criteria. Thiry-four (34.0%) had undergone breast-conserving surgery and 34 had undergone mastectomy. Sentinel node dissection alone had been performed in 73 patients, and axillary lymph node dissection had been performed in 27 patients. Of the 67 patients with hormone receptor-positive breast cancer, 62 (92.5%) received adjuvant hormonal therapy. 53 (53.0%) patients had received adjuvant chemotherapy. Sixty-seven (67.0%) patients had received radio therapy to the breast or chest wall. Recurrence was observed in 14 patients (14%).

**Table. 1.** Clinicopathological characteristics of patients with invasive breast carcinoma of no special type.

| n=100 | No. of patients | % | n=100 | No. of patients | % |
| --- | --- | --- | --- | --- | --- |
| **Age(years) Median(Min,Max)** | 59(27,86) |  | **HER2 status** |  |  |
| **Pathological tumor status** |  |  | negative | 80 | 80% |
| T1 | 56 | 56% | positive | 20 | 20% |
| T2 | 39 | 39% | **Breast surgery** |  |  |
| T3 T4 | 5 | 5% | Mastectomy | 34 | 34% |
| **Pathological lymph node status** |  |  | Breast-conserving surgery | 66 | 66% |
| pN0 | 71 | 71% | **Axillary surgery** |  |  |
| pN1 | 19 | 19% | Sentinel lymph node biopsy | 73 | 73% |
| pN2 | 5 | 5% | Axillary lymph node dissection | 27 | 27% |
| pN3 | 5 | 5% | **Endocrine therapy** |  |  |
| **Histological grade** |  |  | Yes | 62 | 62% |
| Grade 1 | 35 | 35% | No | 38 | 38% |
| Grade 2 | 34 | 34% | **Chemotherapy** |  |  |
| Grade 3 | 31 | 31% | Yes | 53 | 53% |
| **Lymphovascularinvasion** |  |  | No | 47 | 47% |
| Positive | 39 | 39% | **Radiation therapy** |  |  |
| Negative | 61 | 61% | Yes | 67 | 67% |
| **ER status** |  |  | No | 33 | 33% |
| Positive | 67 | 67% | **Recurrence** |  |  |
| Negative | 33 | 33% | Yes | 14 | 14% |
| **PgR status** |  |  | No | 86 | 86% |
| Positive | 49 | 49% |  |  |  |
| Negative | 23 | 23% |  |  |  |
| Unknown | 28 | 28% |  |  |  |

ER, estrogen receptor; PgR, progesterone receptor

Next, we evaluated FOXM1 protein expression by immunohistochemistry. As expected, FOXM1 staining was predominantly observed in the nucleus, although faint cytoplasmic staining was also detected in some cases. We classified FOXM1 protein expression into three scores (0, 1+, 2+, and 3+) based on the percentage of positive tumor cells and performed IHC staining in 22 representative cases. When FOXM1 IHC scores were plotted on the x-axis and *FOXM1* mRNA levels (CPM) on the y-axis, a significant positive correlation was observed between FOXM1 protein expression and its mRNA abundance (p < 0.05; Fig. 1B). This finding supports the consistency between transcriptional and translational levels of FOXM1 expression in invasive ductal carcinoma.

**Figure 1.**
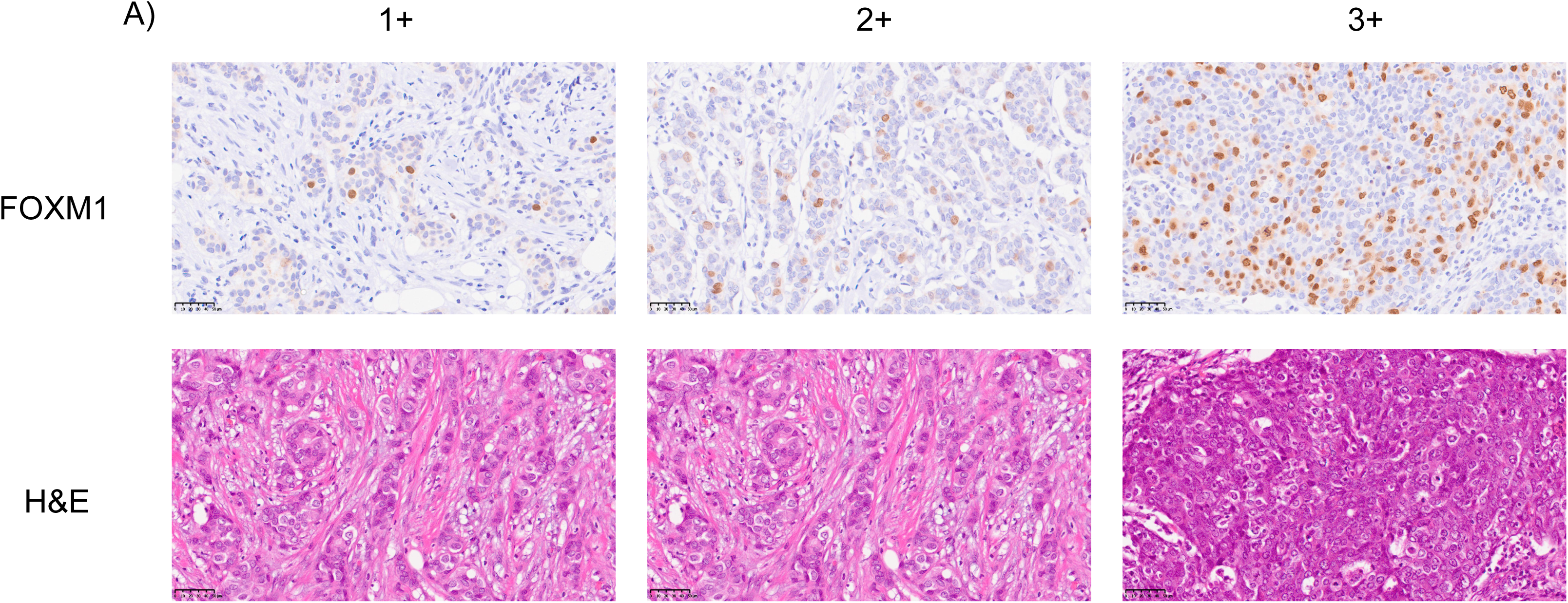
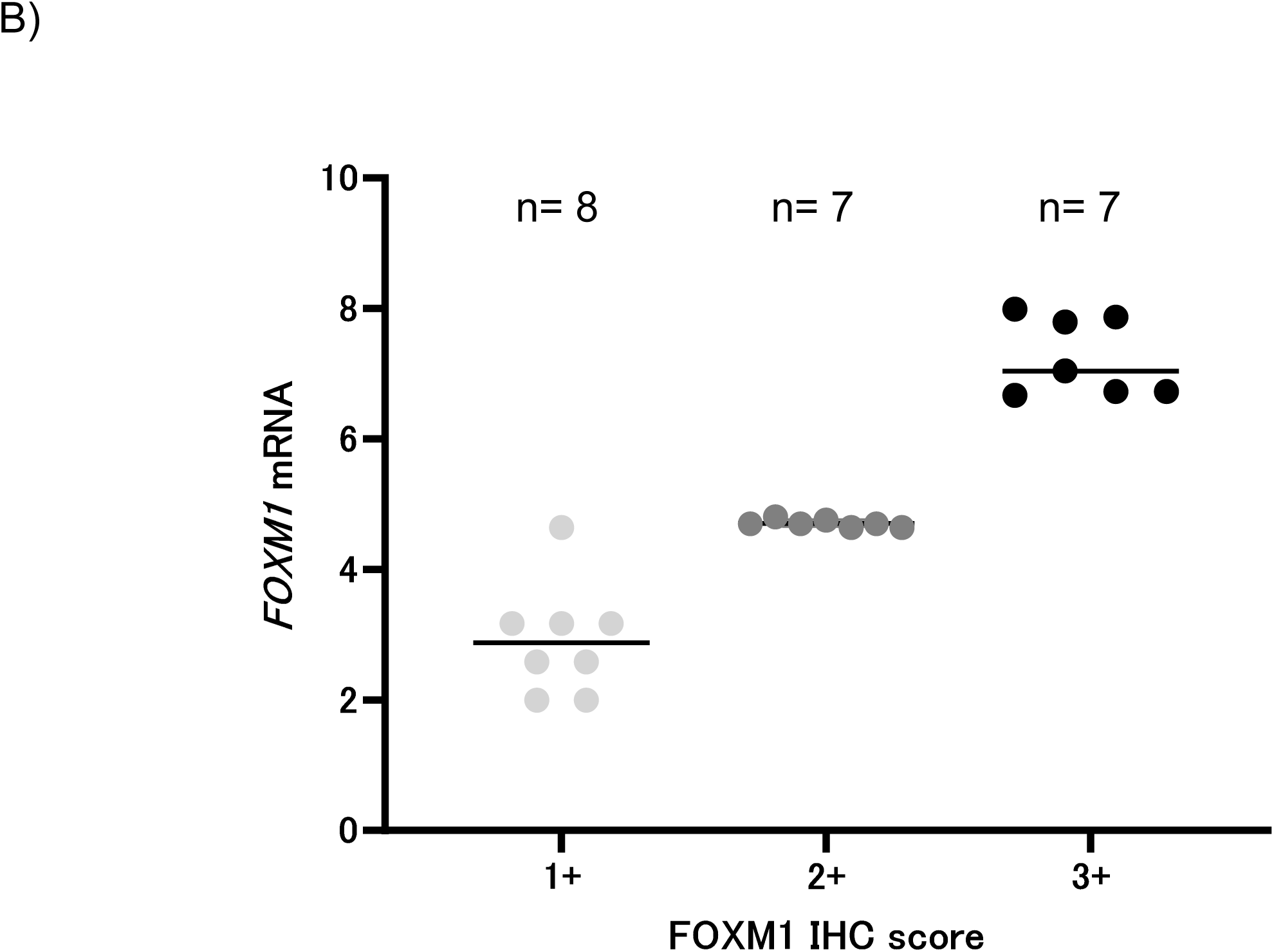
Immunohistochemical analysis of FOXM1 expression in breast cancer tissues and its correlation with mRNA levels. (A) Representative immunohistochemical images of FOXM1 expression scored as 1+, 2+, and 3+, alongside hematoxylin and eosin (H&E) staining of breast cancer tissue sections. Images were captured at 400× magnification. FOXM1 protein expression was evaluated by immunohistochemistry and classified into low, intermediate, and high categories based on staining intensity and distribution. (B) Correlation between FOXM1 protein expression and mRNA levels. FOXM1 IHC scores (x-axis) in 22 representative cases were plotted against *FOXM1* mRNA levels (counts per million, CPM; y-axis), showing a significant positive correlation (p < 0.05).

Receiver operating characteristic curve analysis identified an optimal cut-off value of 4.954 for FOXM1 mRNA CPM to predict recurrence (area under the curve [AUC] = 0.642; sensitivity = 71.4%; specificity = 61.6%; Fig. 2A). At a cut-off value of 4.954, a high *FOXM1* mRNA CPM significantly associated with large tumor, histological III, ER negativity and PgR negativity (Table 2).

**Figure 2.**
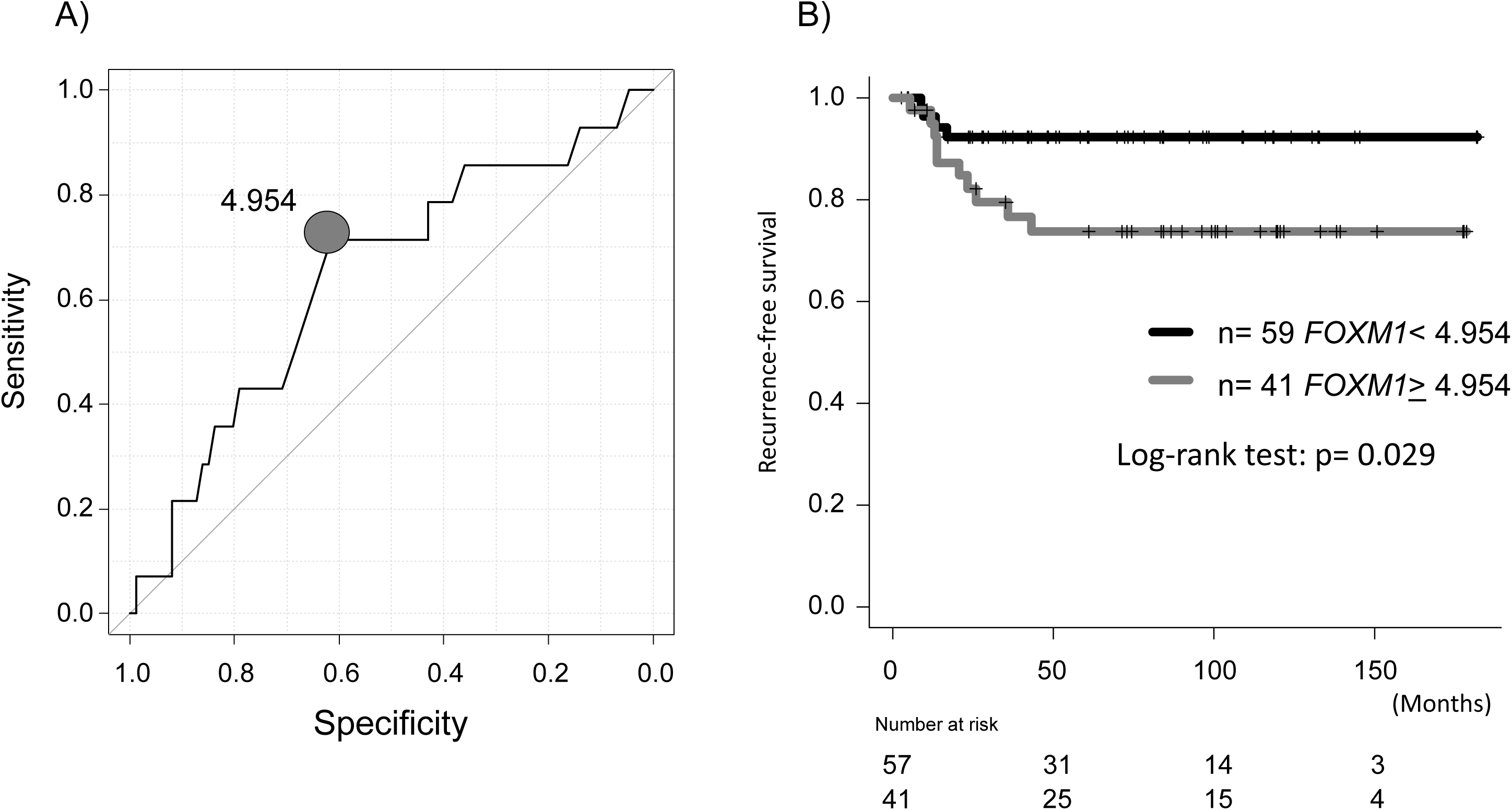
ROC curve and Kaplan–Meier analysis of *FOXM1* mRNA expression and breast cancer recurrence (A) ROC curve identifying the optimal *FOXM1* mRNA CPM cut-off (4.954) for predicting recurrence (AUC = 0.642; sensitivity = 71.4%; specificity = 61.6%). (B) Kaplan–Meier curves showing significantly different 5-year recurrence-free survival between low and high *FOXM1* mRNA expression groups (92.3% vs. 73.6%, p= 0.029).

**Table. 2.** Association between *FOXM1* mRNA CPM expression and clinicopathological parameters in patients with invasive breast carcinoma of no special type.

| Variable |  | FOX M1 mRNA CPM<br>(mean±SD) | P value |
| --- | --- | --- | --- |
| Age | <65 | 4.825 ± 1.142 | 0.171 |
|  | ≥65 | 4.491 ± 1.036 |  |
| pT status | T1 | 4.486 ± 1.079 | 0.015 |
|  | T2 T3 T4 | 5.029 ± 1.101 |  |
| pN status | Negative | 4.638 ± 1.180 | 0.212 |
|  | Positive | 4.949 ± 0.914 |  |
| Histological grade | G1 G2 | 4.265 ± 0.817 | <0.001 |
|  | G3 | 5.749 ± 1.015 |  |
| Lymphovascular |  |  |  |
| invasion | Negative | 4.661 ± 1.149 | 0.474 |
|  | Positive | 4.826 ± 1.072 |  |
| ER status | Positive | 4.349 ± 0.902 | <0.001 |
|  | Negative | 5.488 ± 1.133 |  |
| PgR status | Positive | 4.335 ± 0.908 | <0.001 |
|  | Negative | 5.347 ± 1.374 |  |
| HER2 status | Negative | 4.657 ± 1.216 | 0.226 |
|  | Positive | 4.996 ± 0.505 |  |

CPM, counts per million; SD, standard deviation;
ER, estrogen receptor; PgR, progesterone receptor

Fig. 2B shows a significant difference in 5-year RFS rates between patients with low and high *FOXM1* mRNA CPM (<4.954 vs. >4.954; 92.3% vs. 73.6%, p < 0.001). Univariate analysis of 100 patients identified high FOXM1 mRNA, large tumor size, nodal involvement, and negative hormone receptor status as factors significantly associated with shorter RFS (Table 3). In multivariate analysis adjusting for tumor size, nodal status, nuclear grade, lymphovascular invasion, hormone receptor and HER2 status, and adjuvant treatment, high *FOXM1* mRNA remained an independent prognostic factor (HR 15.26; 95% CI 1.38–169.0; p = 0.026) (Table 3).

**Table. 3.**
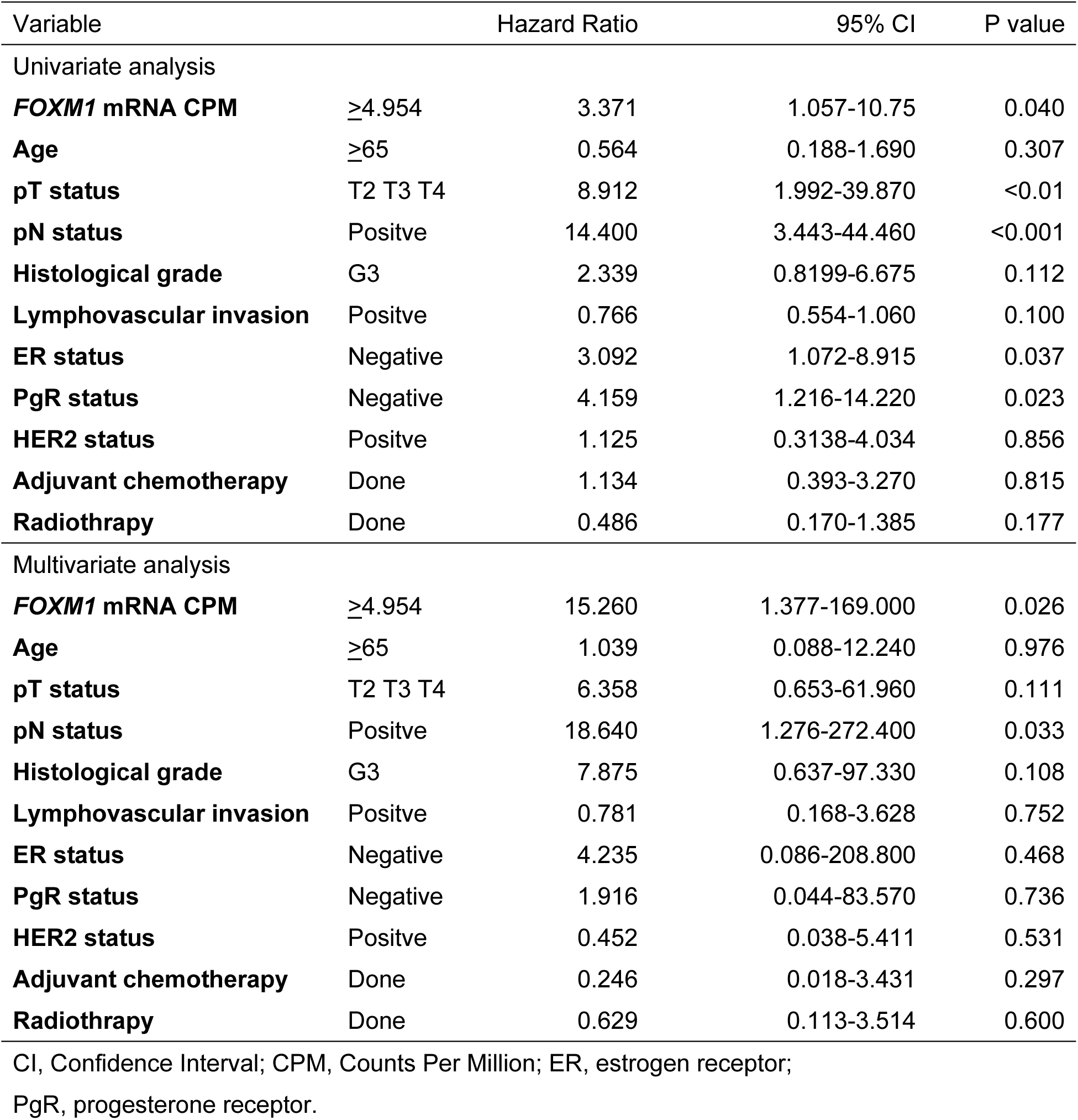
Univariate and multivariate analyses of recurrent-free survival in patients with invasive breast carcinoma of no special type.

Based on RNA-seq data, we compared gene expression profiles between cases with high and low *FOXM1* expression, identifying 190 upregulated and 197 downregulated genes in the high expression group (Fig.3A). Enrichment analyses demonstrated significant activation of cell cycle, chromosome segregation, and mitotic cell cycle processes, as well as pathways related to cellular senescence and motor proteins (Fig.3A). This suggests that high *FOXM1* expression may not only drive proliferation but also contribute to bypassing senescence, thereby promoting tumor progression.

**Figure 3.**
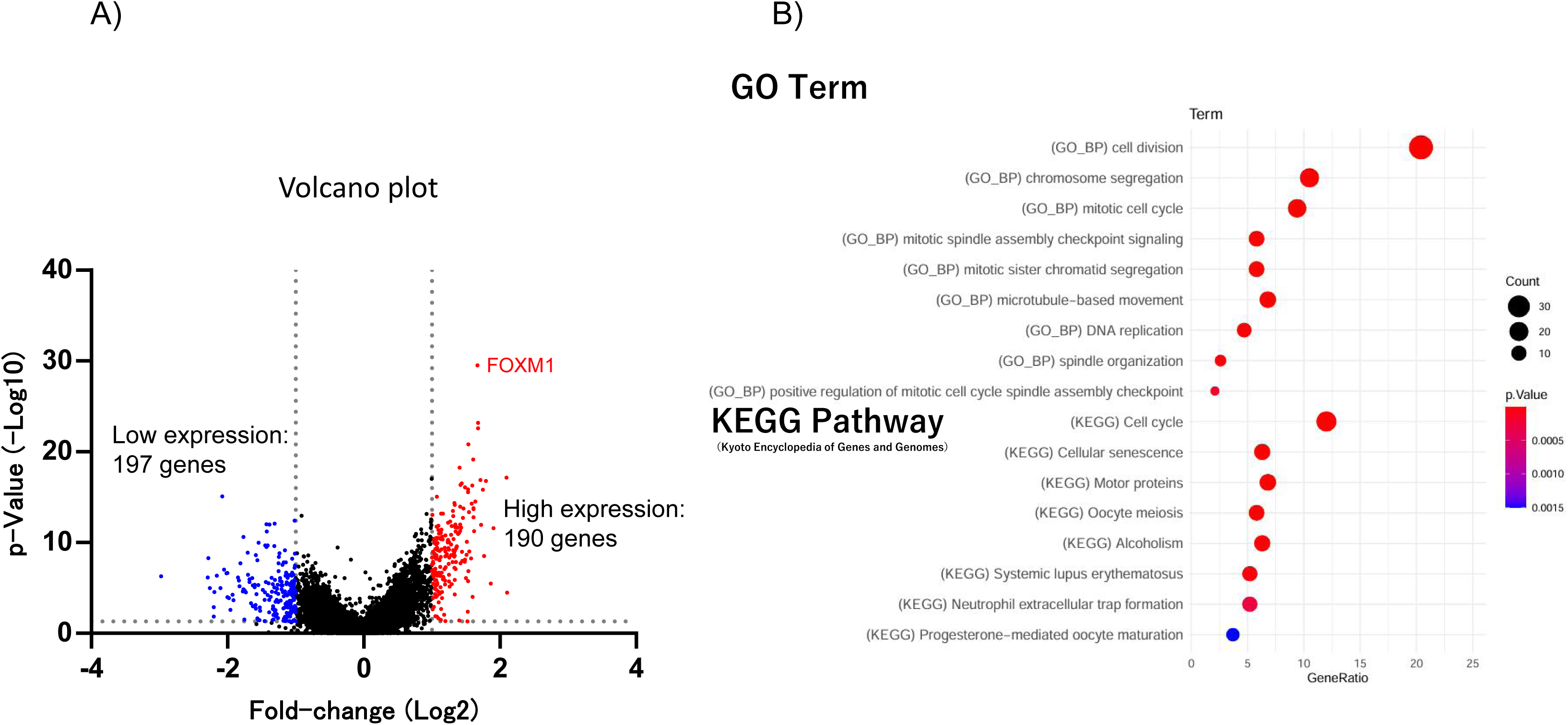
Differential gene expression analysis based on *FOXM1* mRNA CPM cut-off (4.954). (A) Volcano plot illustrating 197 genes with low expression and 190 genes with high expression in relation to *FOXM1* expression levels. CPM, counts per million. (B) Gene Ontology (GO) and Kyoto Encyclopedia of Genes and Genomes (KEGG) pathway enrichment analysis of differentially expressed genes. The top significantly enriched biological processes and pathways are shown for genes upregulated in the high *FOXM1* expression group.

To further dissect the molecular landscape, we constructed a protein–protein interaction (PPI) network centered on FOXM1, revealing ten functional clusters (Fig.4). Notably, FOXM1 occupied a central hub position, connecting genes involved in sister chromatid segregation, DNA replication checkpoint signaling, chromatin structure, and spindle dynamics. Interestingly, the network also included clusters associated with the Ndc80 complex and T-circle formation, raising the possibility of FOXM1 involvement in maintaining chromosomal stability and telomere function. Collectively, these findings underscore FOXM1’s role as a critical regulator that coordinates multiple cell cycle-related and genome integrity pathways in cancer.

**Figure 4.**
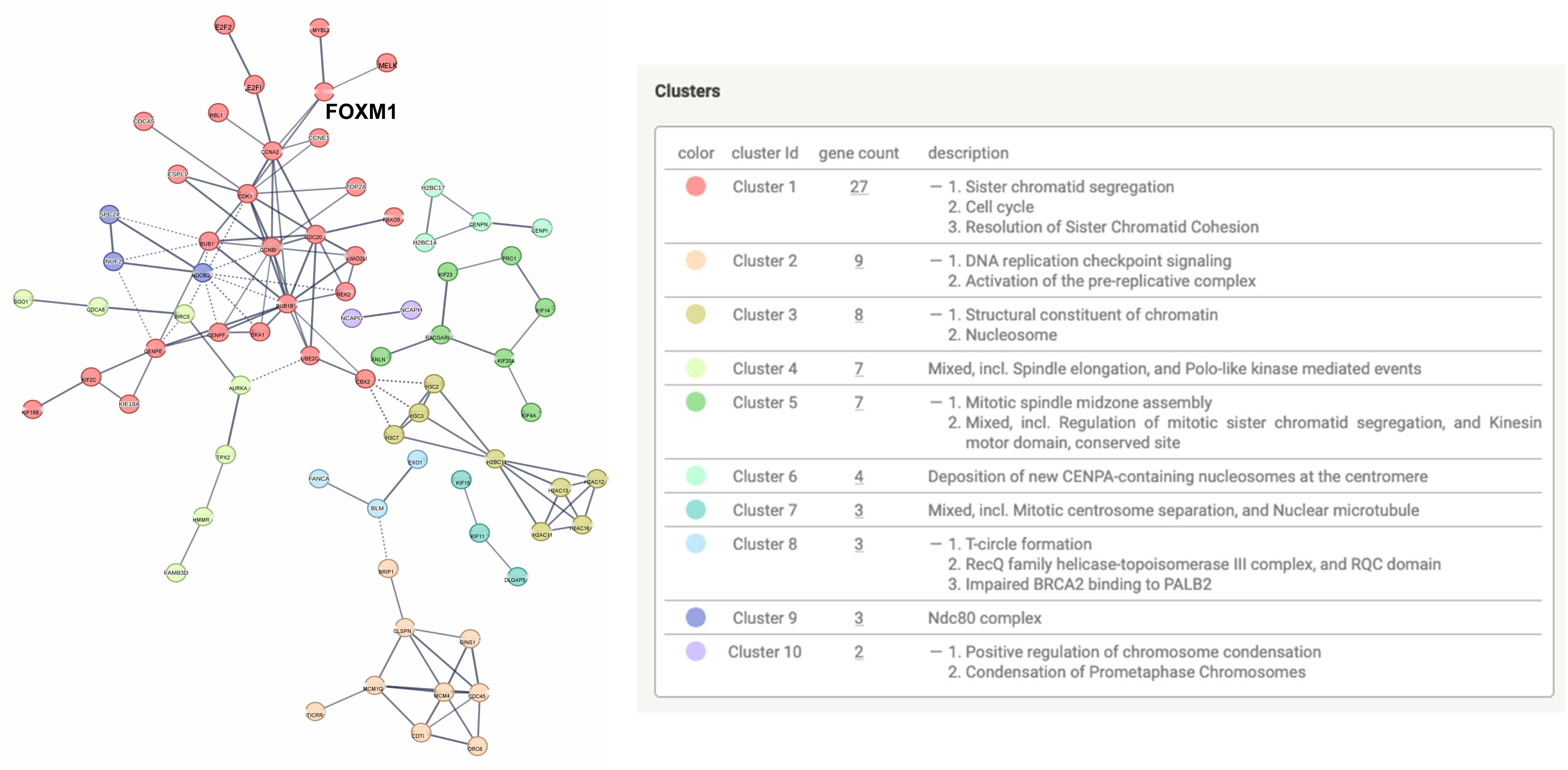
Protein-protein interaction network analysis of genes highly co-expressed with FOXM1. The analysis was performed using the STRING database to identify functional interactions among proteins encoded by genes upregulated alongside FOXM1.

## Discussion

FOXM1 is a key regulator of the cell cycle, playing critical roles at multiple stages, including the G1/S transition, entry into mitosis, and proper progression through mitosis (21). In this study, we identified FOXM1 as a prognostic factor in invasive ductal carcinoma of no special type (IDC-NST), in line with previous reports highlighting its contribution to cancer progression (22,23). Using a cut-off value of 4.954, high FOXM1 mRNA expression was significantly associated with adverse clinicopathological features, including larger tumor size, histological grade III, and negative expression of estrogen and progesterone receptors. Patients with elevated FOXM1 levels also exhibited shorter recurrence-free survival (RFS). Multivariate analysis further confirmed FOXM1 as an independent predictor of poor prognosis, supporting its potential utility as a biomarker for unfavorable outcomes in IDC-NST (24,25). While prior studies have largely assessed FOXM1 by immunohistochemistry, few have validated its prognostic significance at the mRNA level in independent clinical cohorts.

To investigate the mechanisms linking high FOXM1 expression to poor prognosis, we performed gene expression profiling comparing tumors with high versus low FOXM1 levels. Genes associated with cell cycle progression, chromosomal segregation, mitotic spindle organization, and cellular senescence pathways were enriched in the FOXM1-high group, suggesting that overexpression may promote uncontrolled proliferation and help cancer cells evade senescence. These observations align with FOXM1’s role as a transcriptional regulator throughout the cell cycle, whose expression in G1 phase is induced by factors such as c-Myc (26,27). Sequential phosphorylation by cyclin-dependent kinases, PLK1, and other kinases during S, G2, and M phases relieves inhibitory effects on the transactivation domain, thereby activating genes essential for cell cycle progression, including CCNE1, CCNB1, AURKB, CDC25B, and those involved in chromosome segregation and spindle assembly such as CENP-F, KIF20A, and AURKA (28–35). Additionally, FOXM1 promotes proliferation by suppressing nuclear levels of cell cycle inhibitors p21^Cip1 and p27^Kip1 (36) and participates in DNA damage checkpoint regulation via Chk2 phosphorylation (37). Protein–protein interaction analysis further positions FOXM1 as a central hub connecting genes responsible for mitotic checkpoint control, chromatin remodeling, and genomic stability, indicating that its dysregulation may contribute to increased malignancy.

High FOXM1 expression is recognized as a marker of poor prognosis in breast cancer, largely because it confers resistance to endocrine therapy and other anticancer treatments. In ER-positive tumors, FOXM1 can drive tamoxifen resistance by expanding stem-like cell populations and upregulating genes involved in epithelial-mesenchymal transition (EMT), migration, and invasiveness (38,39). It also mediates resistance to HER2-targeted therapies, with inhibition restoring sensitivity to trastuzumab (40). Overexpression can further induce resistance to chemotherapeutic agents such as cisplatin and paclitaxel by altering microtubule dynamics (41,42), and emerging evidence suggests that FOXM1 contributes to CDK4/6 inhibitor resistance through anti-senescence mechanisms, modulated in part by interactions with p53, Rb, and PLK1 (43–45). These mechanisms collectively contribute to therapeutic resistance and adverse outcomes, making FOXM1 a promising target to improve treatment responses (31,38,39,46–48).

Due to its frequent overexpression and association with poor outcomes, FOXM1 has emerged as a potential therapeutic target. Small-molecule inhibitors such as RCM-1 and STL427944 have shown antitumor efficacy in preclinical models with minimal toxicity. Combination strategies with chemotherapeutic agents or CDK4/6 inhibitors demonstrate synergistic effects. Moreover, FOXM1 regulates PD-L1 expression, suggesting that its inhibition could enhance the effectiveness of immune checkpoint blockade (49). Recent studies also indicate that FOXM1 undergoes liquid–liquid phase separation in breast tumor cells, forming transcriptional condensates that promote metastasis. Targeting this process with AMPK agonists or specific peptides has shown potential to reduce tumor growth and improve immunotherapy responses (50).

However, this study has limitations. The sample size was relatively small, limiting statistical power, and the observational design precludes definitive conclusions regarding causality. Additionally, focusing on a single cohort of IDC-NST patients may limit generalizability. Future studies with larger, more diverse populations and longitudinal data are needed to validate FOXM1 as a prognostic and therapeutic marker. Further mechanistic investigations are also warranted to fully understand its role in tumor progression and interactions with other signaling pathways. Finally, the development and clinical evaluation of FOXM1-targeted therapies, alone or in combination with existing treatments, will be essential to determine their impact on patient outcomes.

## Conclusion

Our study demonstrates that FOXM1 overexpression at both the mRNA and protein levels is significantly associated with aggressive clinicopathological features and poor recurrence-free survival in invasive ductal carcinoma of no special type. FOXM1 acts as a central regulator of cell cycle progression, chromosomal stability, and mitotic processes, contributing to tumor proliferation and progression. Importantly, FOXM1 serves as an independent prognostic biomarker, underscoring its potential clinical utility in risk stratification. Given its key role in tumor biology and association with therapeutic resistance, FOXM1 represents a promising target for novel therapeutic strategies. Further studies are warranted to validate these findings in larger cohorts and to develop effective FOXM1-targeted treatments to improve patient outcomes in breast cancer.

## Acknowledgments

We sincerely thank Editage (www.editage.com) for English language editing.

We also appreciate the valuable assistance provided by the clinical research coordinators at Saitama Medical University, Noriko Wakui and Sachiko Aihara, in data acquisition and management.

## Authors’ contributions

MO, EM, WI, TK, AN, AS, YI, AF, AN, AA, HS, HY, KM, HI, NF, AO, and TS provided the clinical data included in the text, performed the literature search, and confirm the authenticity of all the raw data. KA and MH performed the RNA-seq data analysis. TH conducted the pathological evaluation and other related pathological assessments. MO wrote the manuscript draft. KM, AO, HI, and TS contributed to the conceptualization of the work and interpreted and revised the laboratory test results included in this report. MO critically revised the manuscript and modified the text.

All authors read and approved the final manuscript.

## Ethical Approval / Informed Consent

The protocol for this study was approved by the institutional review board of the Saitama Medical University International Medical Center (IRB protocol number: # 2023-105). The requirement for individual informed consent was waived because of the retrospective design of the study. Instead, information about the study was disclosed, and patients were given the opportunity to opt out, in accordance with institutional policy.

